# Degradation intermediates as an indicator of mRNA and cell metabolism

**DOI:** 10.1101/2021.08.29.458069

**Authors:** Yusheng Liu, Yiwei Zhang, Hu Nie, Falong Lu, Jiaqiang Wang

## Abstract

Traditional mRNA degradation rate measurements involves complex experimental design with RNA labeling or transcription blocking together with sampling at multiple timepoints^1,2^. These experimental requirements limit the application of transcriptome-wide mRNA degradation rate analysis mainly in cultured cells, but rarely in *in vivo* samples. Therefore, a direct and simple strategy needs to be developed to study mRNA degradation rate. Here, we defined mRNA degradation intermediates as transcripts where decay is about to occur or has partially occurred in the 3’-untranslated regions after poly(A) tail deadenylation, and found that the proportion of mRNA degradation intermediates is a very simple and convenient indicator for evaluating the degradation rate of mRNA in mouse and human cell lines. In addition, we showed that a higher proportion of mRNA degradation intermediates is correlated with faster cell cycle and higher turnover rate of mouse tissues. Further, we validated that in mouse maturing oocytes where transcription is silent^3,4^, the proportion of mRNA degradation intermediates is positively correlated with the mRNA degradation rate. Together, these results demonstrate that degradation intermediates can function as a good indicator of mRNA, cell, and tissue metabolism, and can be easily assayed by total RNA 3’-end sequencing from a single bulk cell sample without the need for drug treatment or multi-timepoint sampling. This finding is of great potential for studies on mRNA degradation rate at the molecular, cellular, or organic level, including samples or systems that cannot be assayed with previous methods. In addition, further application of the findings into single cells will likely greatly aid the identification and study of rare cells with unique cellular metabolism dynamics such as tissue stem cells and tumor cells.

## Introduction

As the bridge between genes and functional proteins, mRNA metabolism is crucial for the control of gene expression, including synthesis, and degradation. The individual biochemical steps of mRNA degradation have been well studied^1,2^. Half-life has been considered a good indicator of mRNA degradation rate^5,6^, because mRNA degradation profiles typically match a simple exponential decay. There are two main approaches to study the mRNA half-lives at the transcriptome scale. The first approach involves global inhibition of transcription with drug or genetic perturbation, and the half-life can then be determined by the loss of mRNA over time since inhibition^6–8^. The second approach involves metabolic labelling, and the half-life can be determined from either the disappearance of the mRNA or the kinetics of initial labelling^1,2,9,10^. However, both approaches require perturbation of transcription, introducing potential bias in quantification of mRNA half-life. In addition, both methods require a large number of cells for analysis over successive time points, limiting the use to model cell lines that can be easily obtained in bulk and accessible to the exogenous treatments. Furthermore, different methods have been shown to yield very different ranges of mRNA half-life estimation for the same cultured cell lines, with a relatively low correlation between the measured half-lives^5,9,11^. Thus, it is very challenging to accurately analyze mRNA metabolism in a noncell line system, such as in tumor migration and embryonic organogenesis.

With few exceptions, mRNA degradation begins with the deadenylation-linked removal of the RNA poly(A) tail^2,12–14^, which is one of the essential structural components of mRNA^15–17^. Thus, in a population of unsynchronized cells for each gene there must be a stable proportion of mRNA degradation intermediates— transcripts where decay is about to occur or has partially occurred in the 3’-untranslated region (UTR) after poly(A) tail deadenylation, because each point of the mRNA metabolism wave has been covered by the large number of unsynchronized cells. This raises the question of whether the proportion of mRNA degradation intermediates reflect the degradation rate of mRNAs.

Here, we found that different genes indeed have different proportion of degradation intermediates among their mRNA transcripts in mouse NIH 3T3 cells (3T3), mouse embryonic stem cells (mES) and human Hela cells (Hela). Additionally, we found a strong correlation between the proportion of mRNA degradation intermediates and the mRNA degradation rate in both unsynchronized bulk cells and synchronized maturing oocytes. These findings suggest that mRNA degradation intermediates can be used as a simple indicator to measure mRNA degradation rate at the transcriptome scale in any samples that can undergo 3’-adaptor ligation-based high-throughput RNA sequencing, such as PAIso-seq2 and TAIL-seq^18,19^.

## Results

### Mathematic model of mRNA degradation rate and the proportion of degradation intermediates

Typical mRNA molecules are transcribed in the nucleus and polyadenylated immediately following transcription. The poly(A)-tailed mRNAs are then exported to the cytoplasm for translation. For a single mRNA molecule life cycle, there is an adenylated stable transcript phase followed by a deadenylated degradation intermediate phase. It has been known for thirty years that, with few exceptions, bulk mRNA undergo decay initiation by poly(A)-tail shortening^2,12–14^. After poly(A) tail removal, the unprotected 3’ end is attacked and digested by a 3’-5’ exonuclease^16^. Therefore, mRNA degradation intermediates, meaning transcripts where decay is about to occur or has partially occurred in the 3’-UTR after poly(A) tail deadenylation, exist widely in cells (Fig. 1a).

**Fig. 1.**
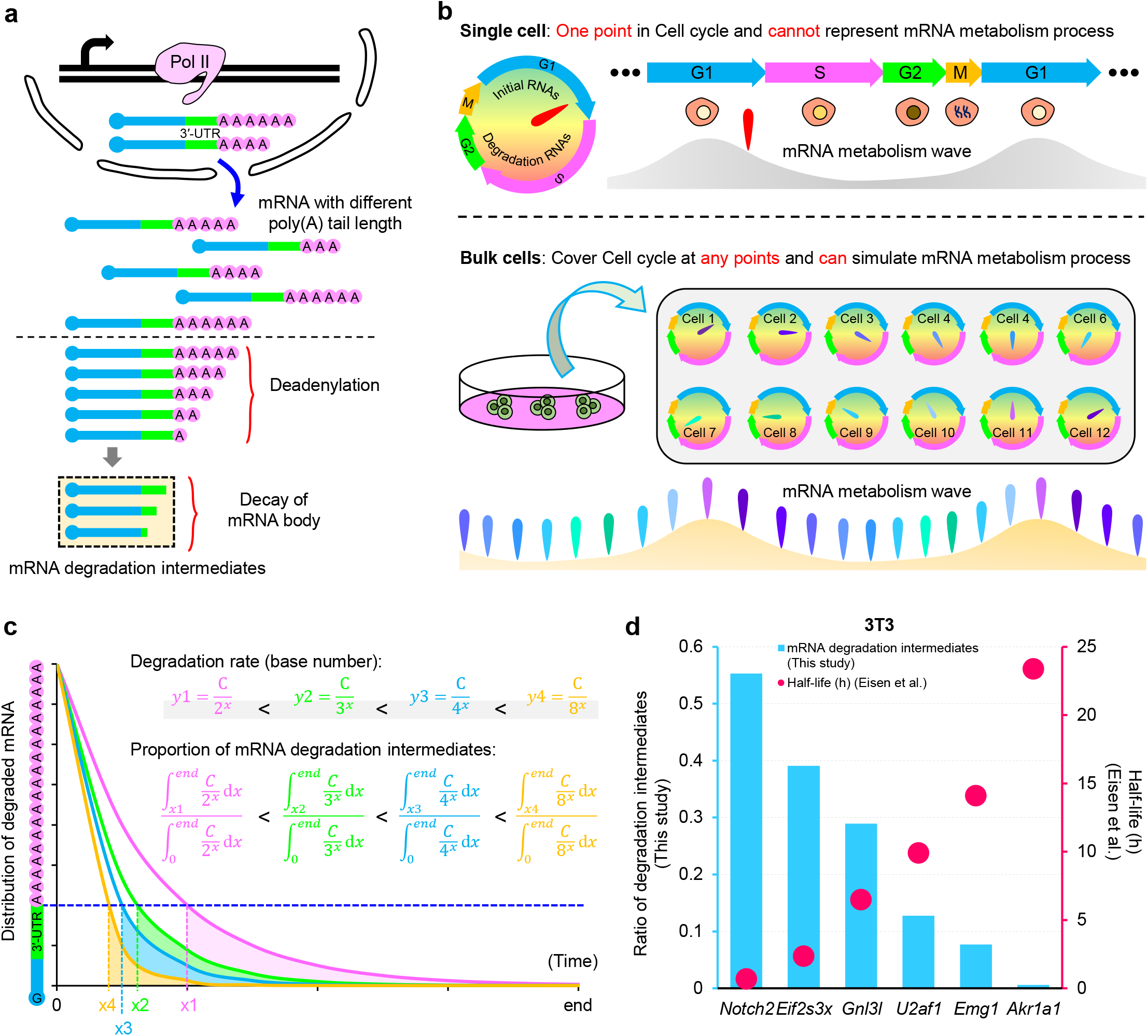
Mathematic model of mRNA metabolism rate and the proportion of mRNA degradation intermediates. **a,** Model showing the generation of tail-less mRNA decay intermediates (mRNA degradation intermediates) in cells. **b,** Model showing that one time point in cell cycle cannot represents mRNA metabolism process in a single cell, while one time point in cell cycle can simulates mRNA metabolism process in bulk cells. **c,** Mathematical model correlating mRNA metabolism rate and the proportion of mRNA degradation intermediates. We put together different exponential decay functions with different base numbers, and normalized them with a same initial length (y = C when x = 0). For function y1, the ratio of the degradation intermediates area (under the y1 line, above the x axis, and right of x1), to the total area (under the y1 line, the x axis, and right to the y axis) represents the proportion of mRNA degradation intermediates. The positive correlation between mRNA metabolism rate and the proportion of mRNA degradation intermediates is shown. **d,** Half-lives measured by data from previous studies^1^ and the proportion of mRNA degradation intermediates measured by PAIso-seq2 for *Notch2*, *Eif2s3x*, *Gnl3l*, *U2af1*, *Emg1*, and *Akr1a1* of 3T3 cells.

In a single cell, mRNA molecules are transcribed, processed, and degraded following an inherent procedure, forming a cell type-specific mRNA metabolism wave as the cell cycle progresses (Fig. 1b, top). Thus, only continuous time points that cover at least one mRNA metabolism wave, but not any single time point, will give an accurate map of mRNA decay for a single cell (Fig. 1b, top). For bulk cells of the same type, in contrast, there are enough number of cells covering any point of the mRNA metabolism wave and any point of the cell cycle at a single time point (Fig. 1b, bottom). Because the proportion of mRNA degradation intermediates can be easily generated from data of any bulk samples, we hypothesize that there should be mathematical models that can 1) depict the whole mRNA metabolism wave of one bulk sample based on data from a single time point, and 2) correlate the proportion of mRNA degradation intermediates with mRNA degradation rate.

Because decay for each transcript typically matches a simple exponential decay function, we built a model using different exponential decay functions with different base numbers, and normalized them with a same initial length (Fig. 1c; y = C when x = 0). We found that the proportion of mRNA degradation intermediates (under the blue dotted line in which the tail has been fully deadenylated) is positively correlated with the base number of exponential decay functions, reflecting the principle that the higher the metabolic rate is, the higher the proportion of mRNA degradation intermediates will be (Fig. 1c). In other words, the longer the mRNA half-life is, the lower the proportion of mRNA degradation intermediates will be.

In order to test our mathematical model, we analyzed half-life data for several representative genes determined by 5-ethynyl uridine (5EU) metabolic labelling^1^ and calculated the proportion of mRNA degradation intermediates from our PAIso-seq2 data of 3T3^19^. There was a negative correlation between half-life and the proportion of mRNA degradation intermediates (Fig. 1d), indicating an increasing proportion of mRNA degradation intermediates along with faster mRNA degradation, which fits our mathematical model. Therefore, we propose that in bulk cells, the proportion of mRNA degradation intermediates is positively correlated with mRNA degradation rate and negatively correlated with mRNA half-life.

### Proportion of mRNA degradation intermediates negatively correlates with mRNA half-life

With our PAIso-seq2 dataset for 3T3 and mES^19^, we calculated the transcriptome-wide proportion of mRNA degradation intermediates and compared it with the mRNA half-life data from prior studies^1,7,8,10^. We collected three mRNA half-life datasets of 3T3 cells: two from multi-time point 5EU metabolic labelling^1^, and the other from single time-point 4-thiouridine (4sU) metabolic labelling^10^. Strong negative correlations (Rp = −0.57 to −0.44) were observed between the proportion of mRNA degradation intermediates and mRNA half-life in all three datasets (Fig. 2a). We also collected three mRNA half-life datasets of mES, all of which use global transcription inhibition by RNA polymerase inhibitor treatment^7,8^. As a result, moderate negative correlations (Rp = −0.37 to −0.25) between the proportion of mRNA degradation intermediates and mRNA half-life were also observed in mES (Fig. 2b).

**Fig. 2.**
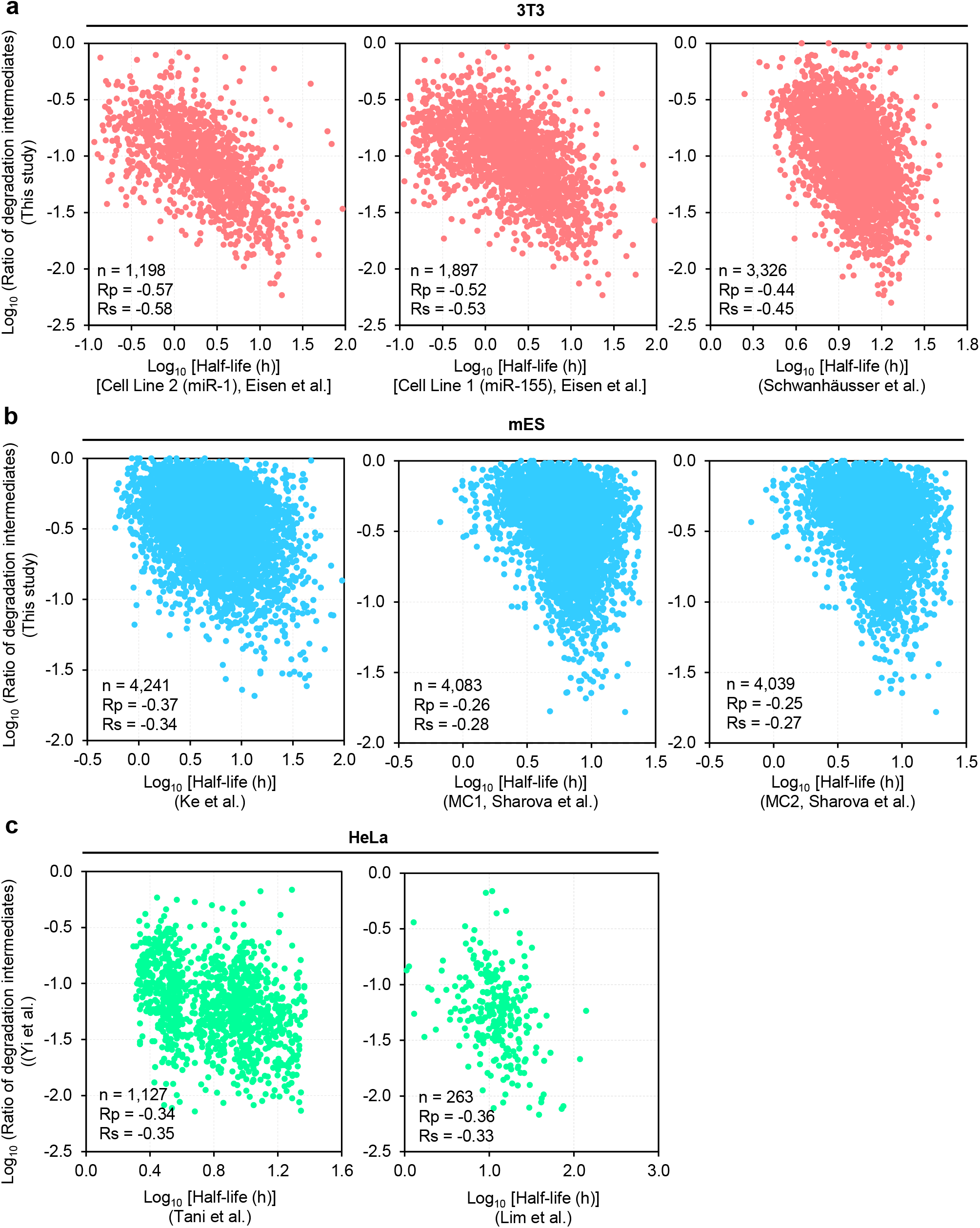
Proportion of mRNA degradation intermediates negatively correlates with mRNA half-life. Scatter plot of the proportion of mRNA degradation intermediates and mRNA half-lives at the transcriptome scale in 3T3^1,10^ (**a**), mES^7,8^ (**b**), and HeLa^20–22^ (**c**) measured by PAIso-seq2. Each dot represents one gene. Genes with at least 10 reads in each sample are included in the analysis. Pearson’s correlation coefficient (Rp), Spearman’s rank correlation coefficient (Rs), and number of genes included in the analysis are shown on the bottom left of each graph.

In addition to the PAIso-seq2 data, we analyzed a published TAIL-seq dataset for HeLa^20,21^, and found that the proportion of mRNA degradation intermediates calculated from TAIL-seq data^22^ showed a negative correlation (Rp = −0.36 to −0.34) with mRNA halflife in HeLa^20,21^ (Fig. 2c). Taken together, these transcriptome-wide analyses reveal that the proportion of mRNA degradation intermediates negatively correlates with mRNA halflife.

### Proportion of mRNA degradation intermediates behaves well to indicate cell cycle duration

The RNA metabolism wave is associated with the cell cycle^23^. In addition, the average/median mRNA half-life is known to be associated with cell cycle duration. For example, the mRNA degradation lifetime is roughly five minutes in *E. coli*, 20 minutes in yeast and 10 hours in human cells; these values are positively correlated with their cell cycle times (roughly 30 minutes for *E. coli*, 90 minutes for yeast, and 50 hours for human cells). Therefore, we speculated that the global proportion of mRNA degradation intermediates would be elevated in rapidly cycling cells (Fig. 3a). To validate this, we compared the proportion of mRNA degradation intermediates at the transcriptome level in the mouse cell lines 3T3 (cell cycle duration: 20 – 26 hours [http://www.nih3t3.com/]) and mES (cell cycle duration: 12 – 17 hours^24^). As expected, the global proportion of mRNA degradation intermediates was significantly higher in mES compared to 3T3 (Fig. 3b). Moreover, the proportion of mRNA degradation intermediates for individual genes in mES were significantly higher than those in 3T3 (Fig. 3c). For example, *Aco2*, *Ybx1*, *Eef1b2*, *Slc25a3*, *Srsf3*, and *Atrx* all showed a higher proportion of mRNA degradation intermediates in mES than in 3T3 (Fig. 3d). Together, these results indicate that the proportion of mRNA degradation intermediates negatively correlate with cell cycle length, making it a good indicator of cell cycle duration time.

**Fig. 3.**
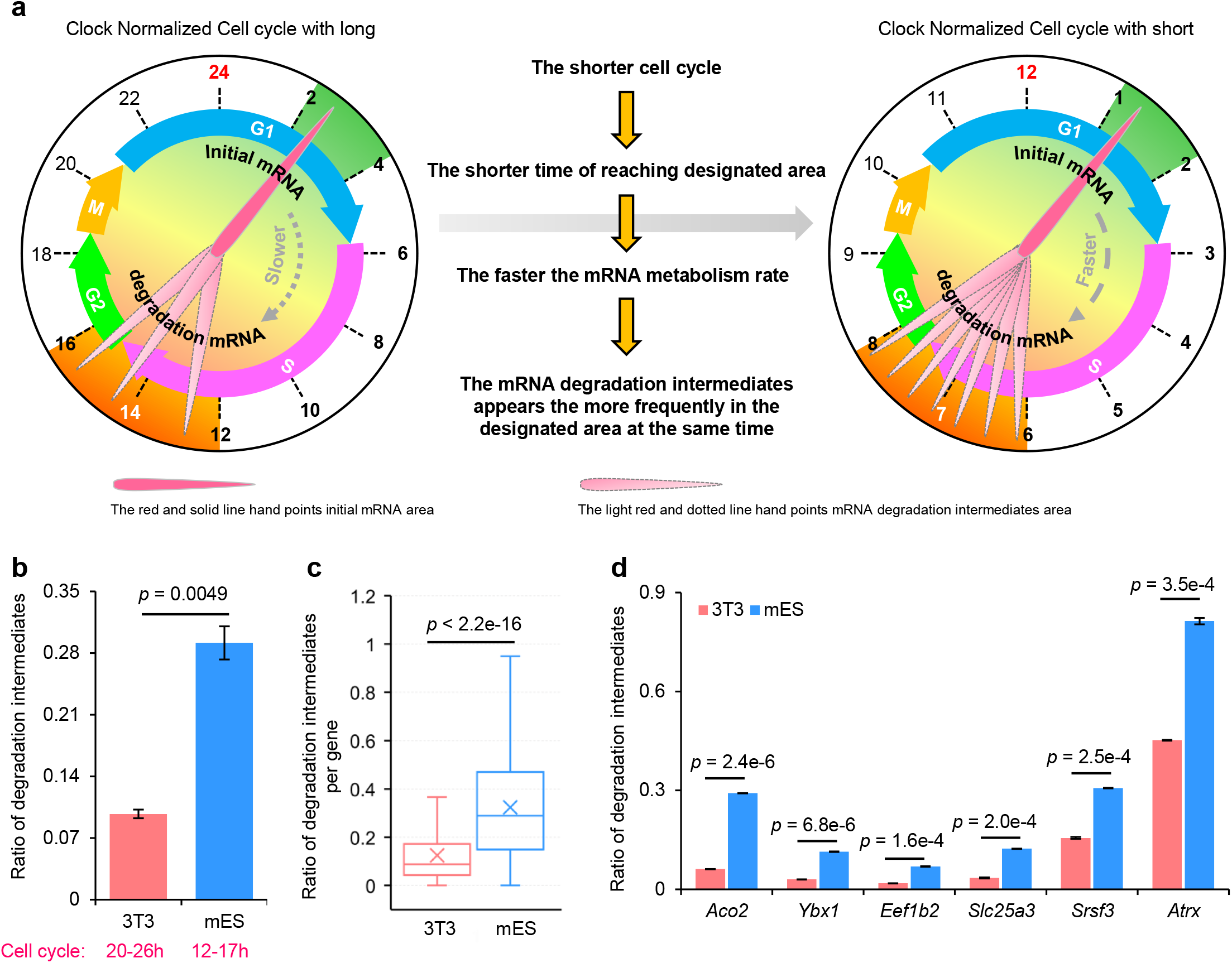
Proportion of mRNA degradation intermediates indicates cell cycle duration. **a,** Model showing that the proportion of mRNA degradation intermediates will be elevated in rapid cycling cells. For different cell cycles, the mRNA metabolism wave is associated with the cell cycle, so the proportion of initial (newly transcribed) mRNAs (dark green) and mRNA degradation intermediates (orange) are similar. For a fixed period of time (for example, 72 h), the hand points to the mRNA degradation intermediates region fewer times in the slower cell cycle (about three times; left) and more times in the faster cell cycle (about six times; right). **b,** Proportion of mRNA degradation intermediates at the transcriptome level in 3T3 and mES measured by PAIso-seq2. Cell cycle durations are shown below the x-axis. Error bars indicate standard error of the mean (SEM) from two replicates. *p-*value was calculated with Student’s *t*-test. **c,** Box plot of the proportion of mRNA degradation intermediates of individual genes in 3T3 and mES measured by PAIso-seq2. Genes with at least 20 reads in the 3T3 and mES sample are included in the analysis (n = 4,348). The “×” indicates the mean value, the horizontal bars show the median value, and the top and bottom of the box represent the value of 25th and 75th percentile, respectively. *p-*value was calculated with Student’s *t*-test. **d,** Proportion of mRNA degradation intermediates of *Aco2*, *Ybx1*, *Eef1b2*, *Slc25a3*, *Srsf3*, and *Atrx* in 3T3 and mES measured by PAIso-seq2. Error bars indicate SEM from two replicates. *p-*value was calculated with Student’s *t*-test.

### Proportion of mRNA degradation intermediates works well as indicator of the mouse tissue turnover rate

To determine whether mRNA degradation can be used to infer the turnover rate of tissues, we calculated the proportion of mRNA degradation intermediates for different mouse tissues^19^, and found that different tissues have different proportions of mRNA degradation intermediates (Fig. 4a). Previous data comparing the turnover rate of mouse muscle to liver, and muscle to heart^25,26^ (Fig. 4b), showed that the absolute turnover rate values are very different. However, the relative turnover rate of these three tissues are clear: the heart took about half the time that muscle took for full turnover, while the liver took less than 30% of the time of muscle (Fig. 4b). Thus, among these three tissues, muscle turnover is slowest, liver turnover is fastest, and heart turnover is in the middle. Importantly, the proportion of mRNA degradation intermediates at the transcriptome level correlates well with the tissue turnover rate for these three tissues (Fig. 4a). In addition, the proportion of mRNA degradation intermediates for individual genes were also correlated with tissue turnover rates (Fig. 4c). For example, the proportion of mRNA degradation intermediates for *Eif4g2*, *Ube2d3*, *Gnas*, and *Cnbp* were correlated with the turnover rate of the corresponding tissue (Fig. 4d). Together, these results revealed that the proportion of mRNA degradation intermediates is positively correlated with tissue turnover rate and negatively correlated with tissue turnover time, and can be used as an indicator of tissue turnover rate.

**Fig. 4.**
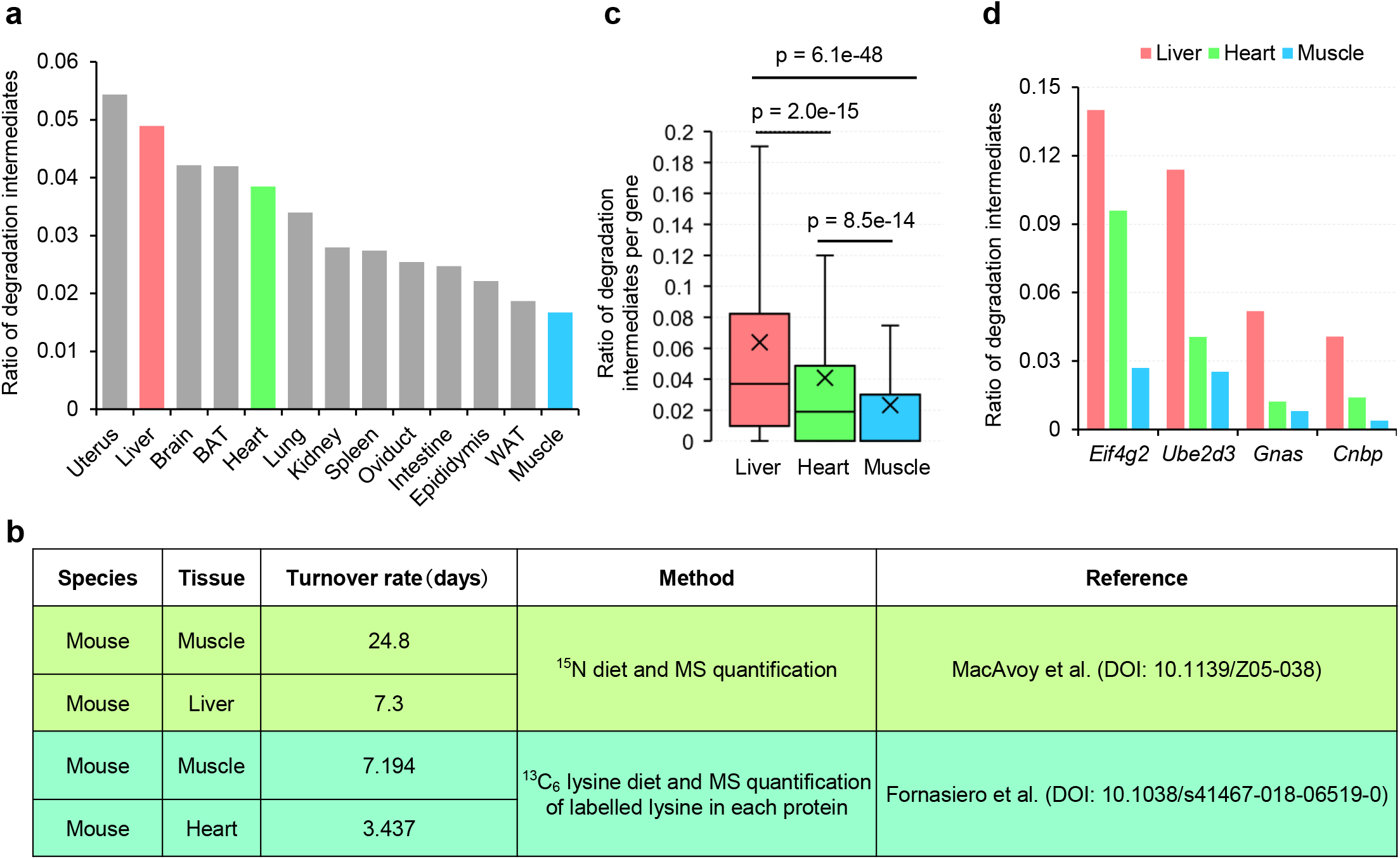
Proportion of mRNA degradation intermediates works well as indicator of the mouse tissue turnover rate. **a,** Proportion of mRNA degradation intermediates across 13 mouse tissues measured by PAIso-seq2. Liver is shown in red; heart is shown in green; muscle is shown in blue; other tissues are shown in gray. **b,** Previously reported data^25,26^ for the turnover time of mouse muscle, liver, and heart tissues. Muscle turnover is the slowest, whereas liver turnover is the fastest, and heart is in the middle. MS, mass spectrum. **c,** Box plot of the proportion of mRNA degradation intermediates of individual genes in mouse liver, heart, and muscle measured by PAIso-seq2. Genes with at least 20 reads in each sample are included in the analysis (n = 1,351, 1,469, and 1,575 for muscle, liver, heart, respectively). The “×” indicates the mean value, the horizontal bars show the median value, and the top and bottom of the box represent the value of 25^th^ and 75^th^ percentile, respectively. *p-*value was calculated with Student’s *t*-test. **d,** Proportion of mRNA degradation intermediates of *Eif4g2*, *Ube2d3*, *Gnas*, and *Cnbp* in mouse liver, heart, and muscle measured by PAIso-seq2.

### Proportion of mRNA degradation intermediates indicates the maternal RNA decay rate during mouse oocyte maturation

Transcription is silent in mammalian fully grown oocytes at germinal vesicle stage (GV oocytes). Therefore, the oocyte maturation process, ranging from GV stage to MI stage and MII stage, provides an ideal physiological system to analyze mRNA decay without the complication of new transcription occurring. We found that the proportion of mRNA degradation intermediates at the transcriptome level increased during oocyte maturation (Fig. 5a), which is consistent with global degradation of mRNA during this process^27^. Consistently, the proportion of mRNA degradation intermediates at the gene level also increased greatly during this process (Fig. 5b). For most of the genes analyzed, the proportion of mRNA degradation intermediates of individual genes showed an increase between different stages during oocyte maturation (Fig. 5c).

**Fig. 5.**
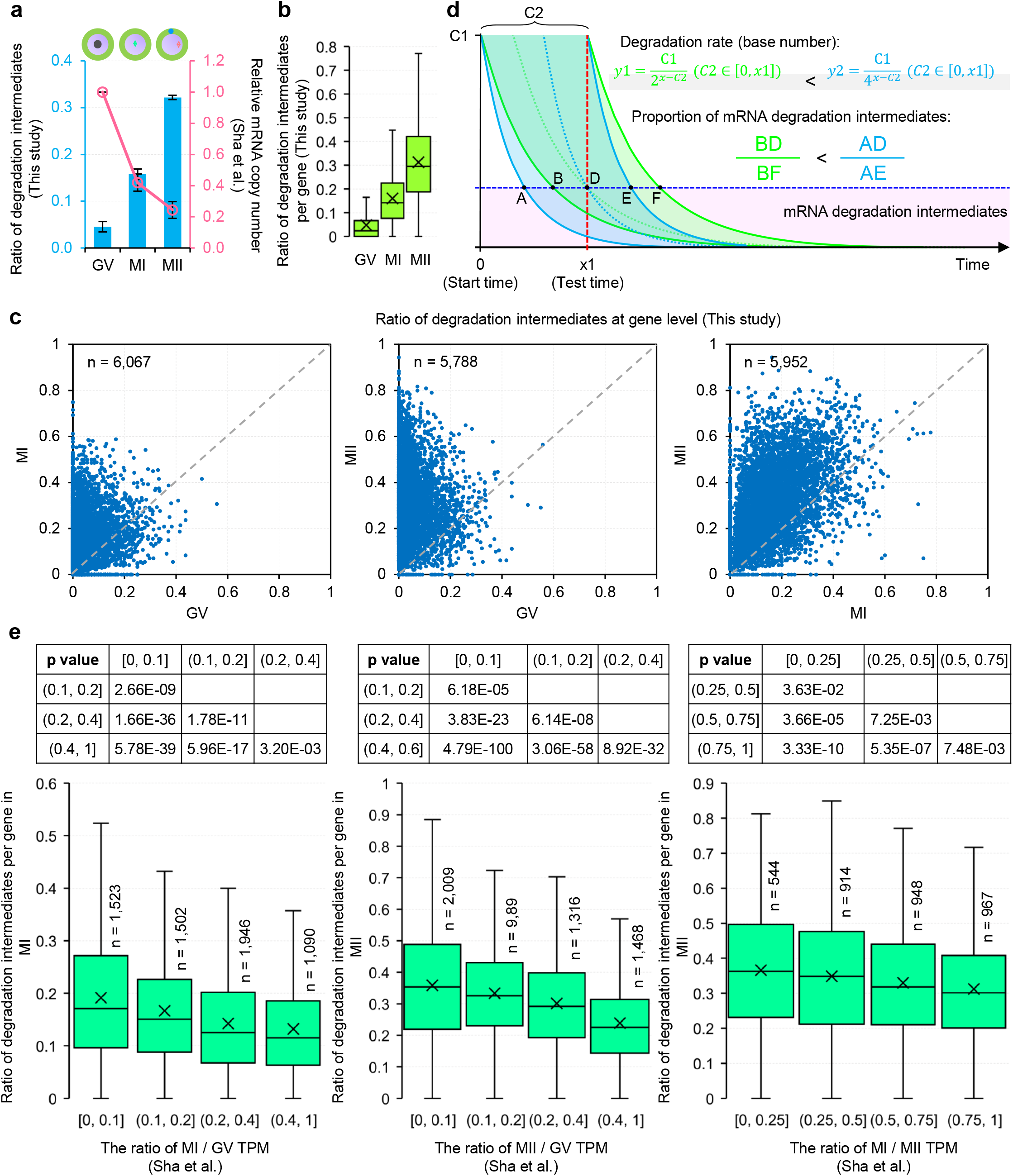
Proportion of mRNA degradation intermediates indicates the maternal mRNA decay rate during mouse oocyte maturation. **a,** Proportion of mRNA degradation intermediates measured by PAIso-seq2 and mRNA abundance (relative mRNA copy number) at the transcriptome level from a previous study^27^ in mouse germinal vesicle (GV), meiosis I (MI), and meiosis II (MII) oocytes. Error bars indicate SEM from two replicates. **b,** Box plot of the proportion of mRNA degradation intermediates of individual genes in mouse GV, MI, and MII oocytes measured by PAIso-seq2. Genes with at least 20 reads in each sample are included in the analysis (n = 6,517, 6,611, and 6,451 for GV, MI, and MII, respectively). **c,** Scatter plot of the proportion of mRNA degradation intermediates of individual genes of samples at neighboring developmental stages in mouse GV, MI, and MII oocytes measured by PAIso-seq2. Each dot represents one gene. Genes with at least 10 reads in each sample are included in the analysis. Number of genes included in the analysis are shown on the top left of each graph. **d,** Mathematical model correlating mRNA degradation rate and the proportion of mRNA degradation intermediates for synchronized maturing oocytes. We put together different exponential decay functions with different base numbers, and normalized them with a same initial length (y = C1 when x = 0). For function y1 and y2, the function translation along the horizontal axis (-C2) represents the continuous degradation of large amounts of mRNAs. At the test time point (x0), for function y1 (y2), ratio of the length of line segment BD (AD) to the length of line segment BF (AE) represents the proportion of mRNA degradation intermediates. The positive correlation between mRNA degradation rate and the proportion of mRNA degradation intermediates is shown. **e,** Box plot of the proportion of mRNA degradation intermediates of individual genes with different mRNA degradation rate during mouse oocyte maturation measured by PAIso-seq2. The mRNA degradation rate is presented by the ratio of TPM of the current stage to the previous stage. Genes with at least 10 reads in each sample were included in the analysis. Number of genes included in the analysis are shown on the top of each box. *p-*value was calculated with Student’s *t*-test. For all box plots, the “×” indicates the mean value, the horizontal bars show the median value, and the top and bottom of the box represent the value of 25^th^ and 75^th^ percentile, respectively.

To determine whether the proportion of mRNA degradation intermediates at the current stage can indicate the mRNA degradation rate from the previous stage to the current stage, we built a mathematical model based on exponential decay functions for synchronized maturing oocytes. The proportion of mRNA degradation intermediates was positively correlated with the base number of exponential decay functions, reflecting that higher degradation rate is correlated with a higher proportion of mRNA degradation intermediates (Fig. 5d). To test the mathematical model, we used the ratio of transcripts per million (TPM) of the current stage to the previous stage as the mRNA degradation rate (smaller value indicates higher decay rate), based on which the genes were grouped into four groups (Fig. 5e). The results showed that mRNA degradation rate was positively correlated with the proportion of mRNA degradation intermediates during oocyte maturation (Fig. 5e). Together, these results reveal that mRNA degradation intermediates can indicate RNA decay rate in both of cell cycle synchronized cells (like maturating oocyte) and unsynchronized bulk cells.

## Discussion

Gene expression is determined by the equilibration of mRNA transcription, and RNA decay^16^. Traditionally, transcriptome-wide mRNA half-lives are determined by RNA quantification after metabolic labelling or global transcription inhibition^1,2^. All these approaches involve drug treatment or genetic perturbation coupled with multiple sampling at a series of time points. These complicated experiments are technically challenging and often yield very different results between different labs for the same cell lines, such as 3T3^1,7,8,10^ (Extended Data Fig. 1). In addition, half-lives are generated through statistical modelling, which is prone to variation caused by different signal to noise levels between experiments. Additionally, drugs that disrupt Pol II-driven transcription also affect stress responses, and other pathways. Thus, not only do the average half-lives differ, but the correlation between half-lives measured with these methods is generally very low^5^. Using unsynchronized bulk cells and synchronized maturing oocytes, we here show that the proportion of mRNA degradation intermediates is a reliable indicator of mRNA degradation rate (Fig. 6). This new indicator has the following advantages: no requirement for drug or genetic treatment; digitalized readout directly from counts, directly comparable between samples and likely between data generated from different labs; and just need a single time point, highly sensitive assay, which enables analysis of low-input samples.

**Fig. 6.**
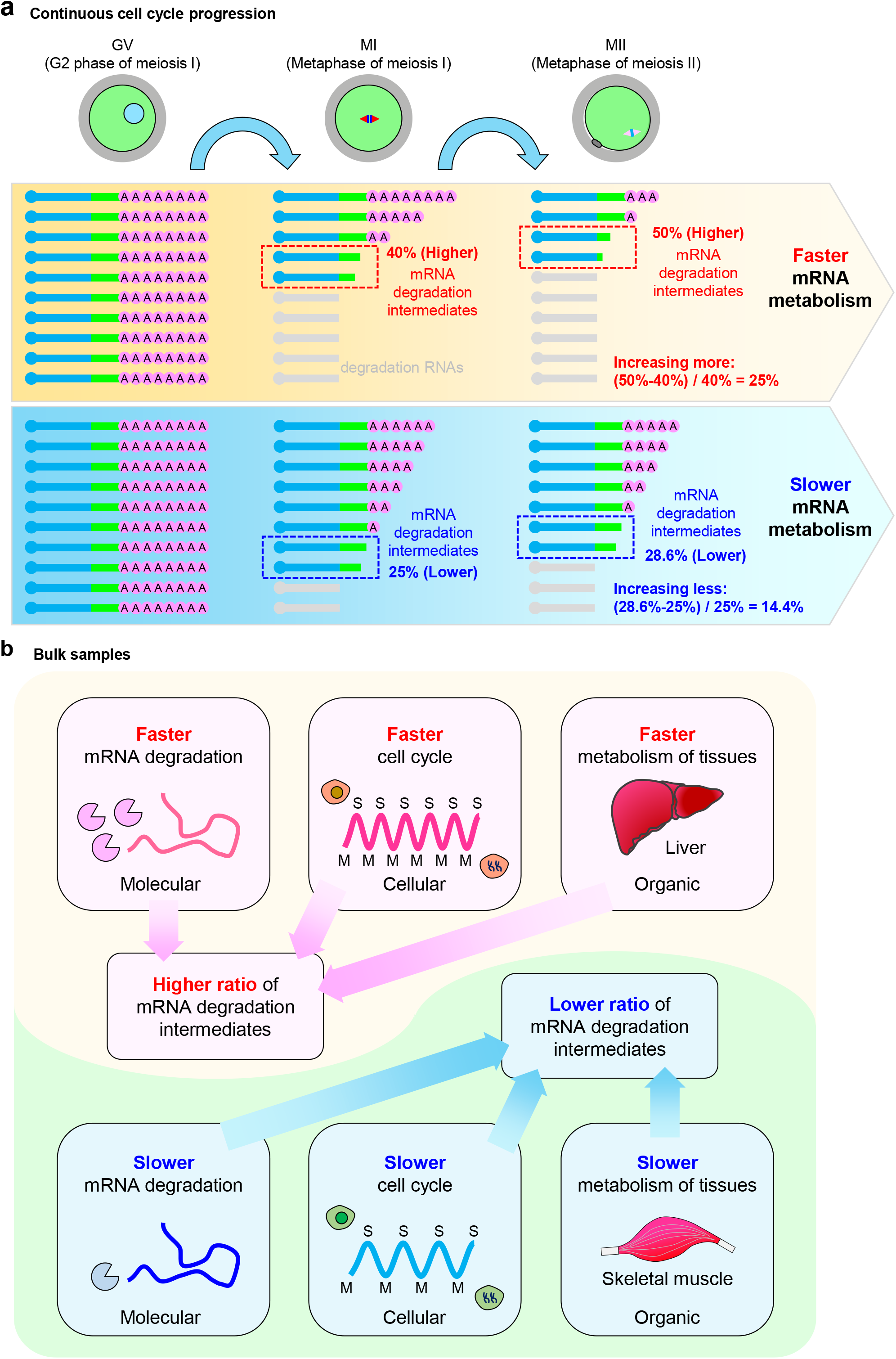
A diagram summarizing mRNA degradation intermediates as an indicator of mRNA, cell, and tissue metabolism in two systems. **a,** Model showing a positive correlation between the proportion of mRNA degradation intermediates and RNA degradation rate during oocyte maturation. **b,** Model showing a positive correlation between the proportion of mRNA degradation intermediates and RNA degradation rate, cell cycle duration, or tissue metabolism rate in bulk samples.

Different cell lines and tissues have different proportions of mRNA degradation intermediates, and higher mRNA metabolism rate of a cell line or tissue corresponds to a higher proportion of mRNA degradation intermediates. In addition, the proportion of mRNA degradation intermediates of the same gene in different tissues also differs. These findings suggest that the proportion of mRNA degradation intermediates is of the potential to distinguish between different cell types. With the continuous progress of single cell technology, more detailed cell typing and functional research are needed. Single-cell mRNA degradation rate measurement maybe can be one of the important indicators for cell typing in the future. However, the current high-throughput single-cell RNA sequencing methods all rely on poly(A) tails to capture mRNAs, which are restricted to polyadenylated mRNA molecules and are completely unable to capture the degradation intermediates. Additionally, current methods cannot evaluate the mRNA degradation rate at the single cell level. Thus, a new single-cell RNA sequencing technology that can sequence the 3’-ends of total RNA regardless of the polyadenylation status (excluding rRNA) would be suitable for mRNA degradation rate evaluation at the single cell level, which will also help to define cell types more accurately.

mRNA metabolism and cell cycle are faster in cancer cells than in normal cells. Thus, we speculate that cancer cells would contain higher proportion of mRNA degradation intermediates than normal cells, which would be a novel indicator for cancer screening of great potential in the future. Additionally, mRNA metabolism may be different in adult stem cells, progenitors, and differentiated cells; thus, the proportion of mRNA degradation intermediates may be useful for identification of adult stem cells, which have key roles in tissue regeneration. We envision that the indicator we have identified here will enable study of mRNA metabolism dynamics during embryonic development as well as disease processes.

## Materials and Methods

### mRNA degradation intermediates analysis

PAIso-seq2 sequencing data were processed as described^19^. Clean CCS reads were aligned to Mus musculus UCSCmm10 reference genome using *minimap2* (v.217-r941)^28^. Then, poly(A) tail was extracted using python script PolyA_trim.py (http://). The 3’-soft clip sequence of the CCS reads in the alignment file is used as candidate poly(A) tail sequence.

The 3’-soft clip sequences with the frequency of U, C and G greater or equal to 0.1 simultaneously were marked as “HIGH_TCG” tails. To better definite the poly(A) tail, we defined a continuous score based on the transitions between the two adjacent nucleotide residues throughout the 3 ‘-soft clip sequences. To calculate continuous score, the transition from one residue to the same residue scored 0, and the transition from one residue to a different residue score 1. The 3’-soft clips which were not marked as “HIGH_TCG” and with continuous score less than or equal to 12 were considered as poly(A) tails. Finally, transcripts without poly(A) tail were treated as mRNA degradation intermediates.

### Statistical analyses

Statistical analyses (mean ± standard error of the mean [SEM]) were performed in Excel. Statistical significance was calculated with Student’s t-test.

## Data availability

PAIso-seq2 data for 3T3 cells, mES cells, 12 mouse tissues (uterus, liver, brain, brown fat, lung, kidney, spleen, oviduct, intestine, epididymis, white fat, and muscle), and mouse oocytes (GV, MI, and MII stages), have been previously described^19,29–31^. The ccs data in bam format from PAIso-seq1 and PAIso-seq2 experiments will be available at Genome Sequence Archive hosted by National Genomic Data Center. This study includes analysis of the following published data: RNA half-life data of Eisen et al. (Gene Expression Omnibus database (GEO) accession no. GSE134660) used in Fig. 1d and Fig. 2a; Schwanhäusser et al. (Sequence Read Archive database (SRA) accession no. SRA030871) used in Fig. 2a; Ke et al. (GEO accession no. GSE86336) used in Fig. 2b; Sharova et al. (GEO accession no. GSE13609) used in Fig. 2b; Tani et al. (DRASearch accession no. DRA000345, DRA000346, DRA000347, DRA000348, and DRA000350) used in Fig. 2c; Lim et al. (GEO accession no. GSE59628) used in Fig. 2c; and mouse oocyte RNA-seq data of Sha, et al. (GEO accession no. GSE118564) used in Fig. 5a. Custom scripts used for data analysis will be available upon request.

## Acknowledgements

This work was supported by the National Key Research and Development Program of China (2018YFA0107001), the Strategic Priority Research Program of the Chinese Academy of Sciences (XDA24020203), National Natural Science Foundation of China (31970588, 32170606), Natural Science Foundation of Heilongjiang province (YQ2020C003), the China Postdoctoral Science Foundation (2020M670516, 2020T130687), and the State Key Laboratory of Molecular Developmental Biology.

## Author Contributions

Yusheng Liu, Falong Lu and Jiaqiang Wang conceived the project and designed the study. Yusheng Liu, Yiwei Zhang, Hu Nie, Falong Lu and Jiaqiang Wang analyzed the sequencing data. Yusheng Liu and Jiaqiang Wang organized all figures. Yusheng Liu, Falong Lu and Jiaqiang Wang supervised the project. Yusheng Liu, Falong Lu and Jiaqiang Wang wrote the manuscript with the input from the other authors.

---

**Extended Data Fig. 1.**
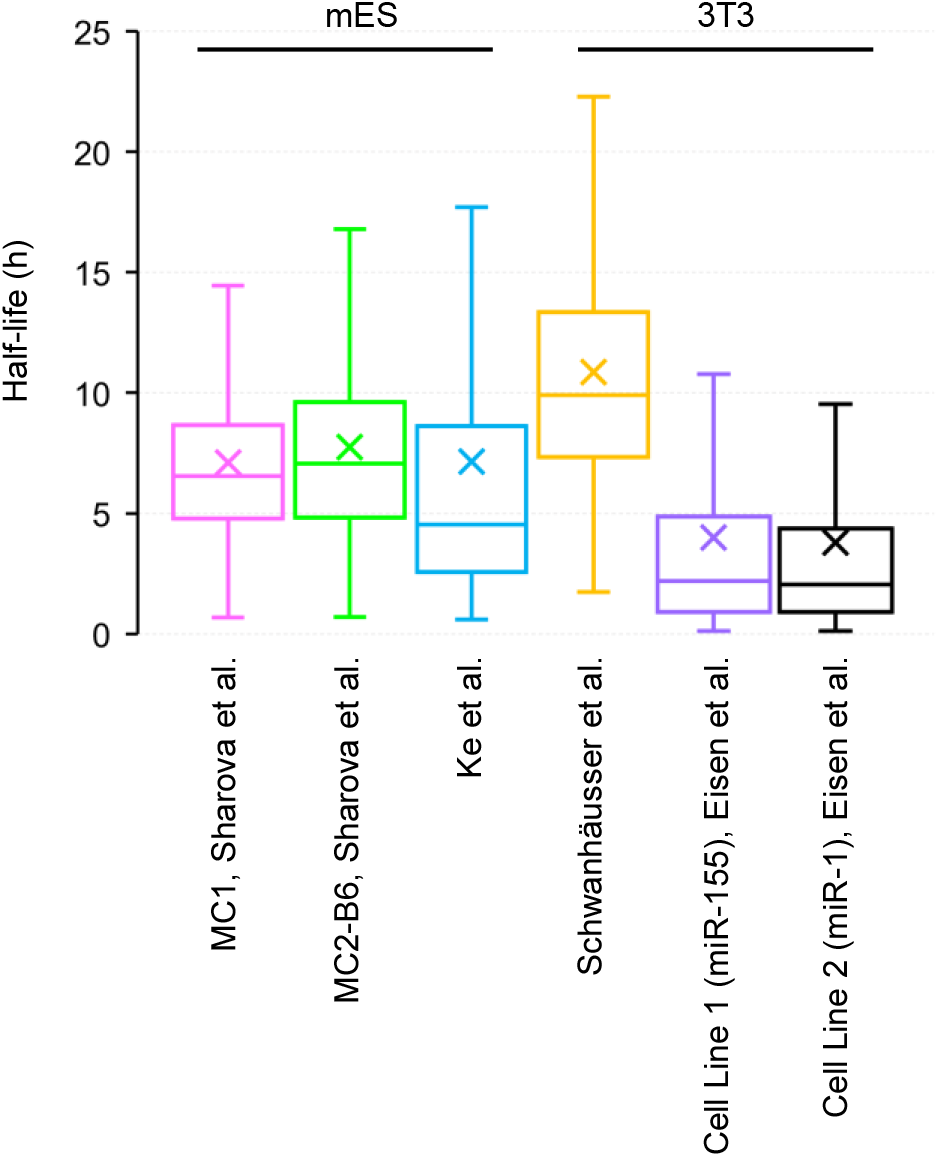
mRNA half-life in 3T3 and mES cells. Box plot of the half-life of mRNA in mES and 3T3 cells from previously reported data^1,7,8,10^.

